# Male recombination produced multiple geographically restricted neo-Y chromosome haplotypes of varying ages that correlate with onset of neo-Y decay in *Drosophila albomicans*

**DOI:** 10.1101/580118

**Authors:** Kevin H-C. Wei, Doris Bachtrog

**Affiliations:** Department of Integrative Biology, University of California Berkeley, CA

## Abstract

Male Drosophila typically have achiasmatic meiosis, and fusions between autosomes and the Y have repeatedly created non-recombining neo-Y chromosomes that degenerate. Intriguingly, *Drosophila nasuta* males recombine, but their close relative *D. albomicans* reverted back to achiasmy after evolving neo-sex chromosomes. Here we use genome-wide polymorphism data to reconstruct the complex evolutionary history of neo-sex chromosomes in *D. albomicans* and examine the effect of recombination and its cessation on the initiation of neo-Y decay. Population and phylogenomic analyses reveal three distinct neo-Y types that are geographically restricted. Due to meiotic exchange with the neo-X, overall nucleotide diversity on the neo-Y is similar to the neo-X but severely reduced within neo-Y types. Consistently, outside of the region proximal to the fusion, the neo-Ys fail to form a monophyletic clade in sliding window trees. Based on tree topology changes, we inferred the recombinant breakpoints that produced haplotypes specific to each neo-Y type and estimated their ages revealing that recombination became suppressed at different time points for the different neo-Y haplotypes. Although there are no evidence of chromosome-wide differentiation between the neo-sex chromosomes, haplotype age correlates with onset of neo-Y decay. Older neo-Y haplotypes show more fixed gene disruption via frameshift indels and down-regulation of neo-Y alleles. Genes are downregulated independently on the different neo-Ys, but are depleted of testes-biased genes across all haplotypes, indicating that genes important for male function are shielded from degeneration. Our results offer a time course of the early progression of Y chromosome evolution, showing how the suppression of recombination, through the reversal to achiasmy in *D. albomicans* males, initiates the process of degeneration.

## INTRODUCTION

Sex chromosomes originate from a pair of homologous autosomes, yet they are often highly differentiated in morphology and function [1]. This transformation starts when a sex-determining factor arises on an autosome causing it to have a sex-specific transmission. Sexually antagonistic mutations, which are variants that are beneficial to one sex but detrimental to the other, are expected to accumulate in close proximity to the sex-determining factor [2,3]. To ensure linkage of the sex-determining factor with sexually antagonistic alleles, recombination between the nascent X and Y chromosomes will become suppressed first around the sex-determining locus. Subsequent gains of sex-beneficial alleles across the chromosome can lead to the expansion of the non-recombining region (for review see [4]).

The cessation of recombination between the sex chromosomes marks a pivotal step that leads the X and Y down a predictable trajectory of differentiation. The absence of recombination on the Y implies that mutations on the same chromosome cannot become unlinked by recombination [2,5,6]. This can lead to the irreversible accumulation of deleterious mutations on the Y, and in the long term, the loss of all of its ancestral genes [1,3,6,7]. In particular, positive selection for an advantageous variant on the Y will cause a sweep of the entire chromosome, dragging along all linked deleterious alleles to fixation [7,8]. Similarly, negative selection on the Y chromosome can greatly reduce its effective population size below its neutral expectation of 1/4 of the effective population size of autosomes, thereby increasing the rate of fixation of weakly deleterious alleles [6,9]. Finally, Y chromosomes may irreversibly accumulate deleterious mutations by a process known as Muller’s ratchet, where Y’s with the fewest number of deleterious mutations are lost by genetic drift [2,10]. All these processes lead to an accumulation of deleterious mutations on the Y, and greatly reduce levels of nucleotide diversity [6,9]. The end result is typically a degenerate chromosome replete with pseudogenes and repetitive elements, and this chromosome is often transcriptionally suppressed by heterochromatin formation [11–16]. During the gradual loss of expression on the Y, the X acquires mechanisms of dosage compensation to balance reduced expression [17]. In Drosophila, this is accomplished by chromosome-wide up-regulation of the X by epigenetic mechanisms [18]. However, it can also be achieved through gene-by-gene up-regulation as exemplified by genes on the Z chromosome of birds [19,20].

Unique to Drosophila and some other insects is the fact that males are achiasmatic, i.e. recombination does not occur during male meiosis. Therefore, autosomes that become fused to the ancestral Y chromosome (so-called neo-Y chromosomes) will be transmitted through males only and will immediately stop any genetic exchange with their former homologs (termed neo-X chromosomes), bypassing the stepwise formation of non-recombining regions. Neo-sex chromosomes have formed independently multiple times in the Drosophila genus by fusions of autosomes (referred to as Muller elements [21,22]) to the ancestral sex chromosomes (Muller A) [23]. Neo-Ys at various state of degeneration within the Drosophila genus reflect their different time of origination. Among the oldest is the neo-Y in *D. pseudoobscura* which is likely the remnant of Muller D after 15 million years of degeneration; the autosomal origin is nearly unrecognizable, with very few genes remaining on its neo-Y [24,25]. Intermediate levels of degeneration can be found on the neo-Y of *D. miranda* which arose ~1.5 million years ago through the fusion of Muller C to the Y [26]. Over 90% of the ancestral protein coding genes can still be identified on the neo-Y, but 40% have become non-functional through frameshifts, nonsense mutations, deletions and transposable element insertions [27–30]. On the younger neo-Y of *D. busckii* which formed through the fusion of Muller F with the sex chromosome around 0.8 million years ago, over 60% of the genes have become pseudogenized [31,32]. Despite the different ages, heavy enrichment of repressive chromatin marks can be found on all these neo-Ys.

One of the youngest known neo-sex chromosome pairs is found in *D. albomicans,* a species primarily distributed across the Asia Pacific island chain (Figure 1) [33,34]. *D. albomicans* is a young species that recently diverged from the west african species *D. nasuta* between 150 and 500 thousand years ago [35,36]. These otherwise indistinguishable sister species produce fully fertile hybrids [37,38], but differ karyotypically by two Robertsonian fusions of the ancestral X and Y with chromosome 3 (henceforth Chr. 3), which is itself a fusion between Muller C and D [39,40]. These two fusions produced two massive neo-sex chromosomes in *D. albomicans* that are each over 55 Mb (Figure 1B). Previous studies have found very few coding differences between the neo-sex chromosomes, but the neo-Y nonetheless shows signs of reduced expression suggesting that degeneration might have begun with disruption of regulatory elements instead of coding sequences [41].

**Figure 1.**
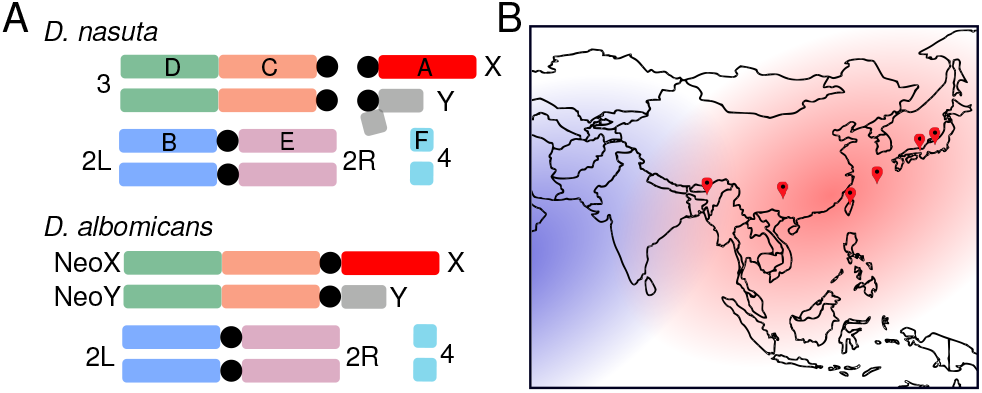
The karyotypes and range of *D. albomicans* and *D. nasuta.* A. The male karyotype of *D. albomicans* and *D. nasuta.* Chromosomes and their corresponding Muller elements are labeled and color-coded. Black circle denotes the centromere. The species ranges of *D. albomicans* and *D. nasuta* are in red and blue, respectively. Red pins denote the sources of the male strains used in this study.

Non-recombining neo-Y chromosomes are expected to harbor low levels of sequence polymorphism [26,42]. Intriguingly, a recent study investigating levels of sequence variability at 27 neosex linked loci found unexpectedly high levels of polymorphism on the neo-Y chromosome of *D. albomicans,* that also has high levels of shared variation with the neo-X chromosome [43]. These findings are inconsistent with either a single origin of the neo-Y, or with recombination being absent between the neo-sex chromosomes. Surprisingly, Satomura et al showed that recombination occurs in *D. nasuta* males, yet is currently absent in *D. albomicans* males [43], suggesting that ancestral recombination between the neo-sex chromosomes in *D. albomicans* males has resulted in the pattern of shared polymorphisms between them.

These surprising observations imply that the young neo-sex chromosomes of *D. albomicans* not only present a window to examine the initiating stages of differentiation, but also provide a unique opportunity to understand how the presence and absence of recombination affects and interacts with the process of Y degeneration. Here, we investigate patterns of neo-Y variation from six population of the species range with whole genome sequencing (Figure 1B). Using population and phylogenetic analyses, we confirm that the neo-Y fusion occurred once, and subsequent meiotic exchange with the neo-X generated multiple neo-Y haplotypes of different ages that are now geographically restricted. Curiously, the Chr. 3 that fused with the ancestral Y carried a *D. nasuta* introgression, which was subsequently broken down by male recombination. We show that different Y-haplotypes have lost recombination at different time points in the past, and their age is associated with different extents of degeneration. These results untangle the complex series of events during the formation and subsequent evolution of the neo-Y in *D. albomicans* and provide timed snapshots of the effect cessation of recombination has on the beginning of sex chromosome differentiation.

## RESULTS

### Complex patterns of evolutionary history on the neo-Y is inconsistent with single origin

In order to characterize the geographic distribution of neo-Y chromosomes, we selected eleven inbred lab strains of *D. albomicans* that were collected from Shilong India, Kunming China, Nankung Taiwan, Okinawa, Kyoto, and Fukui Japan (Supplementary Table 1). The males and females of each strain were genotyped using Illumina high throughput sequencing. Since the current reference is a female assembly lacking the neo-Y chromosome, neo-Y genotypes were determined from neo-X sites that are heterozygous in males but homozygous in females; the neo-Y alleles are then the alleles absent in females. This generated 255,010 variant neo-Y sites across the eleven neo-Ys and 584,333 across the neo-sex chromosomes after filtering. We validated 14 out of 14 neo-Y alleles that coincided with RFLP sites, confirming the efficacy of our approach (Supplementary Table 2). Application of the same bioinformatics pipeline on an autosome identified very few male-specific SNPs confirming that our approach is highly sensitive to detect neo-Y specific SNPs (Supplementary Table 3). Using all sites across the neo-Y chromosome, we constructed a maximum likelihood tree that revealed three distinct groups of neo-Ys (Figure 2A). Group 1 and group 2, henceforth Y1 and Y2, include flies from Taiwan and Japan, with Y2 composed exclusively of flies from the Okinawa Island. The Shilong and Kunming lines form a distinct and distant group, henceforth Y3. The three groups are separated by long branches, but individuals are highly similar within groups. This starkly contrasts to the phylogeny of the neo-X chromosome, where the branches are substantially longer between individuals which lack strong clustering, indicative of higher rates of polymorphisms and less differentiation between populations (Figure 2B).

**Figure 2.**
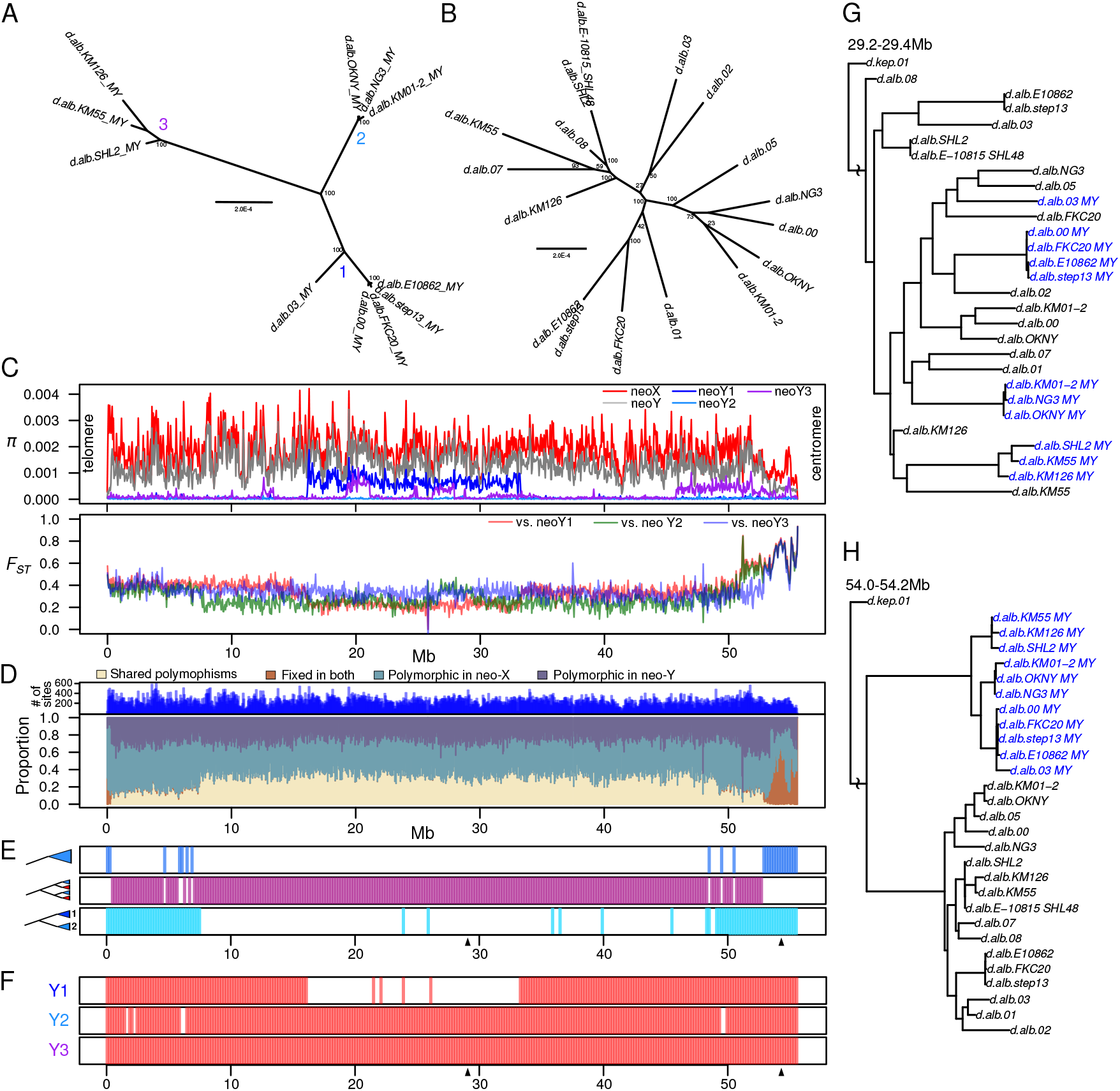
Distinct neo-Y groups due to male recombination. Unrooted maximum likelihood trees for neo-Y (A) and neo-X (B), based on all filtered variant sites across the chromosomes. The neo-Ys are divided into three labeled groups. C. Nucleotide diversity estimates for the neo-X, neo-Y, and neo-Y groups across the chromosome is plotted in the top panel. Population differentiation estimates, F_ST_, between the neo-X and the three Y groups are plotted in the bottom panel. See Supplementary figure 2 for F_ST_ plotted individually. D. The proportion of shared polymorphism and fixed differences between the neo-X and neo-Y across the chromosome is depicted; singletons are removed, and only biallelic sites are used. The number of sites are plotted above. E. Based on 200kb sliding window maximum likelihood trees of the neo-Xs and neo-Ys, three different topologies are plotted: windows where all the neo-Ys form a monophyletic group are plotted in the top track; windows where the neo-Ys are paraphyletic are plotted in the middle track; windows where Y1 and Y2 individuals are monophyletic are plotted in the bottom track. Arrow heads underneath the tracks denote windows where the examples (G-H) are taken from. F. Same as E, but shows windows where individuals within the same Y type are monophyletic. G-H. Trees at windows that exemplify the different topologies; neo-Ys highlighted in blue.

The formation of a Drosophila neo-Y chromosome is typically accompanied with a reduction of effective population size and nucleotide diversity (π). Furthermore, because the fusion that formed the neo-Y likely happened only once and recently [40], π is expected to be minimal due to severe bottlenecking associated with the fusion. While π in the region proximal to the centromere is low, π across the rest of the neo-Y is only marginally lower than π of the neo-X (Figure 2C). Interestingly, within each Y group, we find large swaths of the chromosomes with minimal π, indicating that the Ys within a group are nearly identical (Figure 2C). Therefore, the elevated π is the result of highly differentiated neo-Y chromosomes groups, as further evidenced by high levels of F_ST_ between the groups (Supplementary Figure 1). For all the Y groups, differentiation from the neo-X is extremely high with F_ST_ > 0.6 in regions proximal to the centromere, followed by precipitous drops to ~0.4 (Figure 2C, Supplementary Figures 2-3). For Y1 and Y2, the drops occur more distally from the centromere at 50.5 Mb than for Y3 at 53Mb. Y1 and Y2 both see additional valleys of F_ST_ spanning large parts of the chromosome arm. The drops in F_ST_ are primarily due to increases in shared polymorphism between the neo-X and neo-Y along the chromosome (Figure 2D). The centromere proximal region and the telomeric end are the only region with a high proportion of fixed differences (Figure 2D).

To better understand the causes of these variable patterns of differentiation, we generated maximum likelihood trees for the neo-Xs and neo-Ys in 200kb sliding windows across the chromosome. We find that the centromere proximal region shows the expected topology, where all the neo-Ys form a monophyletic clade with a deep split from all the neo-Xs, further affirming that the neo-Y fusion occurred once shortly after the species split from *D. nasuta* (Figures 2E,H). However, monophyly is interrupted at 53.2Mb which coincides with the decline in differentiation between Y3 and the neo-X (Figure 2C). For the vast majority of the chromosomal windows, the neo-Ys are interspersed within the neo-Xs, instead of forming a monophyletic clade (Figures 2E,G). Y1 and Y2 group together for more regions along the neo-sex chromosome (Figure 2E); the regions where they are no longer monophyletic are associated with reduced F_ST_ and increased levels of shared polymorphism with the neo-Xs (Figures 2C-D). Notably, even though the three Y groups are paraphyletic for most of the chromosome, individuals within them are always grouped together (Figure 2F). The only exception is the taiwanese *D. albomicans* (d.alb 03) which is paraphyletic to the rest of the Y1 individuals for trees in the middle of the chromosome (Figure 2F,G). This topology change is also reflected by elevated π within Y1 and a valley of F_ST_ between Y1 and the neo-X (Figures 2C-D). Despite supporting the single origin of the neo-Y fusion near the centromere, the complex patterns of differentiation and tree topologies across the rest of the chromosome are unlikely to result from a single origin. Instead, these results are consistent with multiple haplotypes of neo-Ys that have been introduced through recombination with the neo-X.

### Male recombination produced large neo-Y haplotype blocks of different ages

The valleys of Fst and plateaus of shared polymorphism between the neo-X and neoY suggest that the different neo-Ys have megabase-sized haplotype blocks. To determine the recombination breakpoints that generated the different haplotypes, we reconstructed the individuals’ haplotypes based on the tree topologies across the chromosome, reasoning that disruption of monophyly of the neo-Ys are due to recombination events with neo-Xs (Figure 3A). For example, monophyly of the neo-Y at the centromere proximal region indicates that there is only one haplotype; this haplotype is disrupted when Y3 is no longer monophyletic with the rest of the neo-Ys, indicating a second haplotype. We find that each Y group has an unique haplotype resulting from recombination breakpoints distal to the centromere (Figure 3B). The largest haplotype block spans nearly the entirety of the chromosome from 0.4 Mb to 53.6 Mb and distinguished Y3 from the others. A slightly smaller haplotype block between 7.6Mb and 49.6Mb distinguishes Y2 from Y1. While we initially categorized the Taiwanese strain (d.alb.03) as Y1, it harbors a different haplotype in the middle of the chromosome separating it from the rest of the Y1 individuals; this also explains the elevated π within the group (Figure 1C). Interestingly, all the individuals have the same haplotype at both ends of the chromosome. While the centromeric end is expected due to the single origin of the Y-autosome fusion, the monophyly of the telomeric regions is surprising, and may suggest selection near the telomere leading to the fixation of a particular haplotype variant.

**Figure 3.**
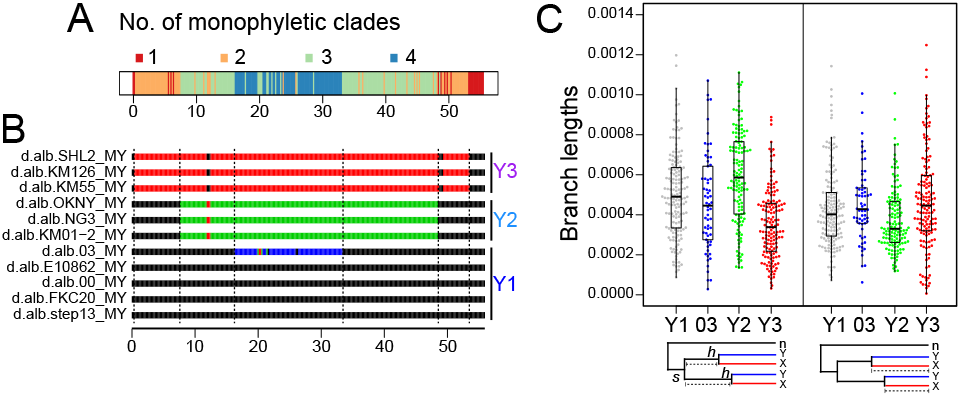
Neo-Y haplotypes and their ages. A. The number of monophyletic branches to which the neo-Ys exclusively belong are inferred from the sliding window phylogenies and plotted. B. For each 200kb window of each neo-Y individual, those that belong to the same monophyletic branch, and therefore haplotypes, are color-coded the same. When single 200kb windows interrupt haplotype blocks, their colors are changed to match the flanking windows. Note the colors here are arbitrary and do not indicate the ancestral haplotype; e.g. the blue segment in d.alb03 could have resulted from recombination in the either the d.alb03 lineage or lineage ancestral to the rest of the Y1s. The breakpoints of haplotypes are demarcated by dotted lines. C. The branch lengths between *s* (node of species split) and *h’s* (nodes of neo-X and neo-Y haplotype splits) are plotted for each haplotype on the left. Shorter lengths indicate older haplotypes. The branch lengths between h’s and their respective neo-X tips are plotted on the right and longer lengths indicate older haplotypes.

Given that *D. albomicans* males no longer recombine [43], we reasoned that since the neo-Y haplotypes would be locked after the cessation of recombination, the number of nucleotide changes before and after can be used to roughly date the age of the haplotypes. We first took the branch length from the node of the species split (s) to the nodes that separate the neo-Y groups from the neo-X sisters (h) (Table 1, Figure 3C left). Shorter branch length indicates that the neo-Y haplotype ceased to recombine with the neo-X sooner after the formation of the species, and, extensively, an older neo-Y haplotype. The Y3 and Y2 haplotypes have the shortest and longest branch lengths, respectively.

**Table 1.** Age neo-Y haplotypes

|  | Median branch length |  | Proportion of *h*<br>to tip | Age (Kya)* |
| --- | --- | --- | --- | --- |
|  | *s* to *h* | *h* to tip |  |  |
| Y1 | 0.000490 | 0.000402 | 0.4564 | 114.1 |
| Y1 03 | 0.000445 | 0.000428 | 0.5064 | 126.6 |
| Y2 | 0.000586 | 0.000330 | 0.3792 | 94.8 |
| Y3 | 0.000338 | 0.000446 | 0.5483 | 137.1 |

Second, we calculated the branch lengths from the nodes separating the neo-Y and neo-X to the tip of the trees, which indicate the amount of changes after the cessation of recombination. However, since the neo-Y haplotypes likely began to experience very different rates of change after recombination stopped, we took the branch lengths of the neo-X sisters instead (Table 1, Figure 3C right). Consistent with the first method, Y3 has the longest branch length and Y2 the shortest. These results indicate that Y3 and Y2 are the oldest and youngest haplotypes, respectively, with Y1 and the taiwanese haplotypes being the intermediates. Based on the the ratio of the two branch length measures and the estimated species age of ~250 Kya, we estimate that the Y1 haplotype stopped recombining roughly 110 Kya, and Y3 and Y2 are roughly 20 Kya older and younger respectively (Table 1). The different ages of the haplotypes and time of cessation of male recombination may indicate the sequential introduction of the causal allele to suppress male recombination in the different populations.

### Fixed *D. nasuta* introgression on the neo-Y is broken down by male recombination

To better understand the evolutionary history of the neo-sex chromosomes after their formation from an autosome, we compared them to *D. nasuta’s* Chr. 3. With the addition of nine *D. nasuta* strains, we inferred 742,084 variant sites between the neo-sex chromosomes and Chr.3. Based on a model whereby the neo-X and neo-Y fusions occurred after the species split, the two chromosomes are expected to be more similar to each other than to Chr. 3. However, lower levels of differentiation between the neo-X and Chr. 3 can result from gene flow between the two species since they recombine in female hybrids. Due to this and slightly lowered π, the neo-Y shows higher levels of F_ST_ from Chr. 3 than the neo-X does for nearly the entire length of the chromosome. However, at the centromere proximal region, we see a surprising reversal where the neo-X is more differentiated from Chr. 3 (Figure 4A); this region, as mentioned earlier, also shows sharply elevated Fst between the neo-X and all neo-Y groups (Figure 2C and 4A). The absolute divergence (D_XY_) between neo-X and Chr. 3 is also significantly higher at this region than the D_XY_ between neo-Y and Chr. 3, even though the values are similar for the rest of the chromosome (Figures 4A-B). These results indicate that the centromere proximal region of the neo-Y is more similar to Chr. 3 than to the neo-X, inconsistent with a simple model whereby the neo-Y fusion occurred after the species split.

**Figure 4.**
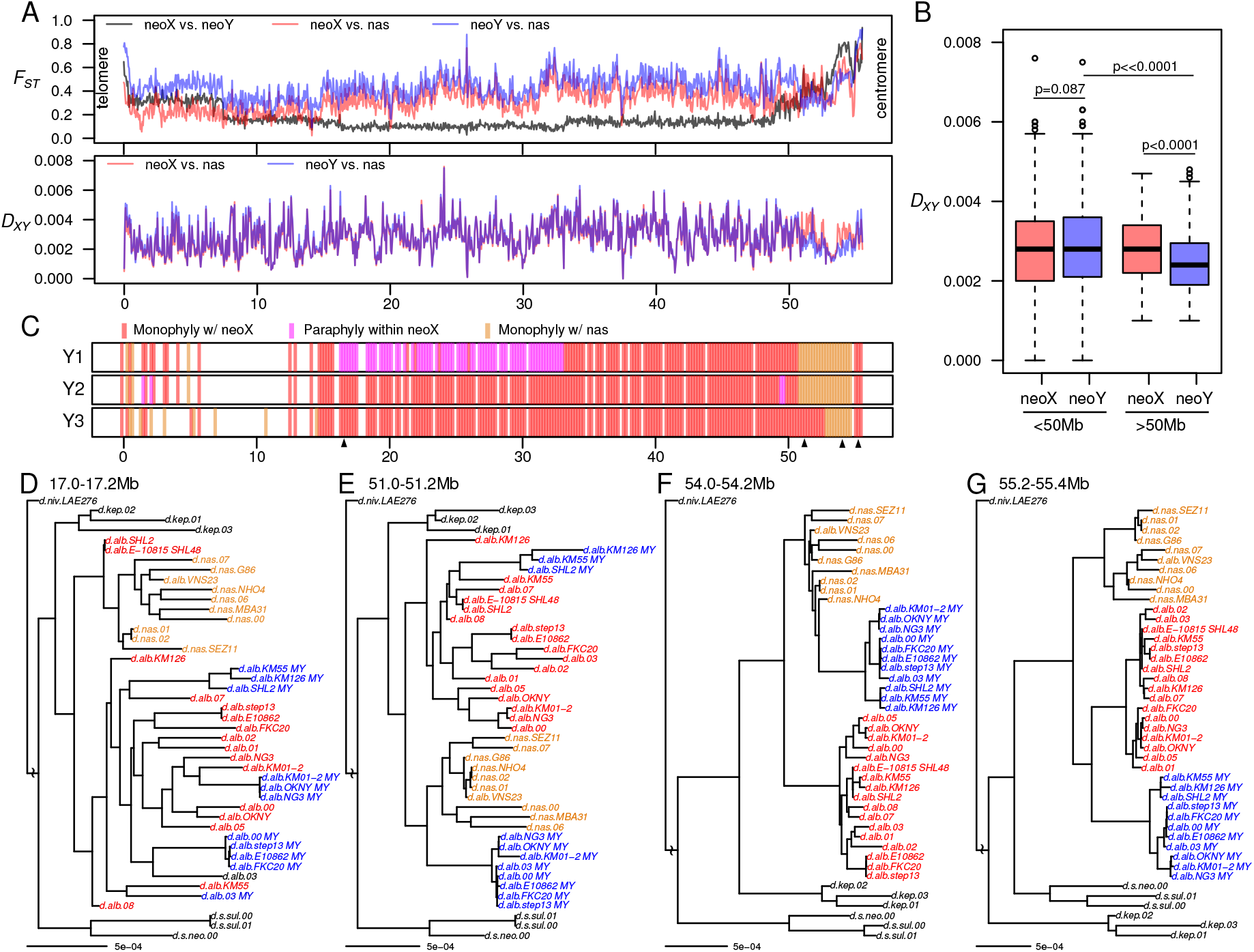
*D. nasuta* introgression on the neo-Y. A. The *F_ST_* between the neo-sex chromosomes (gray), the neo-X and *D. nasuta* Chr. 3 (red), and the neo-Y and Chr. 3 (blue) are plotted in the top panel. The absolute divergence *(D_XY_)* of neo-X (red) and neo-Y (blue) versus Chr. 3 are in the lower panel. B. The distribution of *D_XY_* (from 4A) from 0-50Mb and 50-55Mb are shown in boxplots, with the p-values above the relevant comparisons (Wilcoxon’s rank sum test). C. Based on sliding-window 200kb maximum likelihood trees of neo-Xs, neo-Ys, and Chr. 3s, the phylogenetic relationships between each Y1 group to the neo-X and Chr. 3s are determined. Windows in which the neo-Ys fall within or are sisters to neo-Xs either monophyletically or paraphyletically are plotted in red or magenta, respectively. Windows in which they fall within or are sisters to Chr. 3 are marked yellow. Missing windows are those in which the neo-Xs and Chr. 3s do not sort cleanly into separate clades, likely due to introgression or incomplete lineage sorting. The SHL2 and SHL48 lines are removed from the phylogenies prior to topology inference. D-G Representative windows (demarcated by arrowheads in 4C) of notable topologies are displayed, with the neo-Xs, neo-Ys and Chr. 3s colored in red, blue, and yellow, respectively.

We constructed maximum likelihood trees with the neo-Xs, neo-Ys, and Chr. 3s along the chromosome to decipher the unexpected similarity between the neo-Y and Chr. 3. While the neo-Ys are monophyletic to the neo-Xs from 53Mb to the end of the chromosome (Figure 2D), we find two different topologies when incorporating Chr. 3s. For ~0.6 Mb directly next to the centromere, the monophyletic neo-Y clade is sister to the neo-X (Figure 4C and G). The topology then changes such that the neo-Y clade clusters with/within the *D. nasuta* clade (Figure 4C and F), revealing that the centromere proximal region of the neo-Y contains an introgression. The introgressed region is found within all three Y groups, and thus most likely fixed on the neo-Y. This introgression is roughly 4.2 Mb in Y1 and Y2 and 2.0 Mb shorter on Y3. The distal end of Y3’s shorter introgression block lands precisely at a haplotype breakpoint (Figure 4C and E), arguing that male recombination truncated the introgression with sequences from the neo-X. Given that the introgression is fixed, we suspect that the neo-Y fusion happened between the ancestral Y and a recombinant Chr. 3, instead of recombination after the fusion that introduced the *D. nasuta* introgression. Unlike the latter scenario which not only requires male recombination, but also subsequent fixation of the introgression, in the former scenario, the recombinant chromosome can be produced by female hybrids. Supporting this possibility, we find that the neo-X of one of our *D. albomicans* lines (d.alb.SHL2) contains a large introgression from *D. nasuta* (Figure 4D and Supplementary figure 2). This Indian line was collected in the western edge of the *D. albomicans* range overlapping with the *D. nasuta* range. While no hybrids have been reported or collected in the wild, this line and our results clearly indicate the importance of hybridization in the process of speciation.

### No evidence of chromosome-wide down-regulation

In the absence of male recombination, the neo-Y is expected to start accumulating deleterious mutations immediately after its formation. Indeed, genes on older neo-Y chromosomes have been found to be down-regulated due to pseudogenization and spreading of heterochromatin from adjacent repeats [6,30,44]. The neo-X, in turn, will become dosage compensated through transcriptional up-regulation, in order to balance the down-regulation of the neo-Y homologs [45,46]. Previously, Zhou and Bachtrog reported that the neo-sex chromosomes in *D. albomicans* show widespread down-regulation of neo-Y linked genes, despite few pseudogenes; 30.1% of the genes were found to show significant neo-X bias and the neo-X alleles are on average ~1.3-fold more highly expressed than the neo-Y counterparts [41]. However, our reanalysis of the RNA-seq data of the KM55 strain, belonging to Y3, reveals little difference between the expression of the neo-X and neo-Y alleles with a median fold-difference of 1.04, a negligible bias that is also observed in the DNA sequence (Figure 5A, Table 2). Out of 3181 genes with distinguishable neo-sex chromosome alleles, only 115 (3.6%) show significant neo-X bias. Similarly we see no evidence for chromosome wide reduction in expression for Y1 (d.alb.03) and Y2 (d.alb.OKNY-2) (Figure 5A, Table 2), with few significantly neo-X biased genes and negligible median expression difference between the alleles. The reason for this difference are not entirely clear, and may reflect differences in mapping strategies, reference allele mapping bias, and assembly quality.

**Figure 5.**
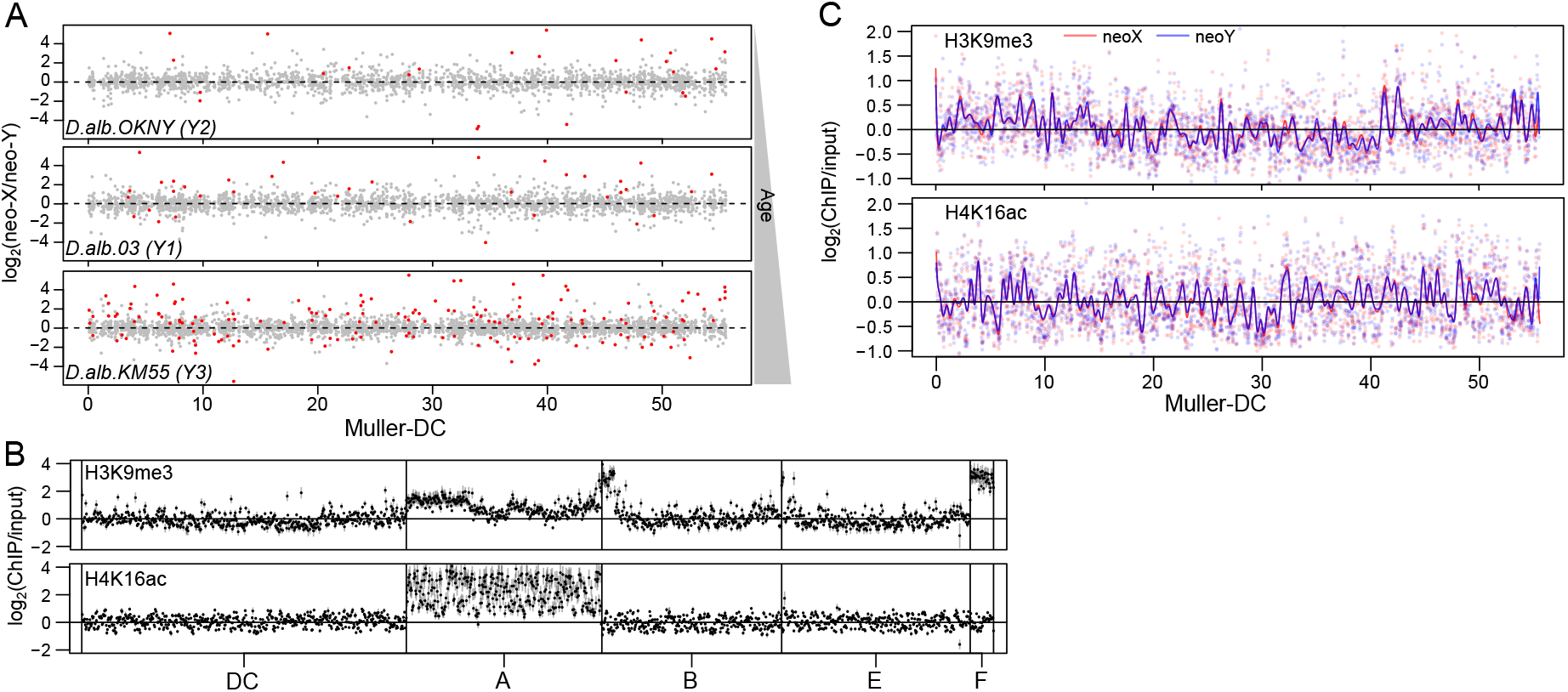
Absence of chromosome-wide differentiation. A. Based on allele-specific RNA-seq, the corrected fold-difference (see materials and methods) between the expression of neo-X and neo-Y alleles of each gene is plotted for neo-Ys of different ages: OKNY (Y2), 03 (Y1), and KM55 (Y3). B. Enrichment of the repressive (H3K9me3, top) and dosage compensation (H4K16ac, bottom) marks for strain 03 are plotted in 50kb windows genomewide; error bars represents the standard errors inferred from quadruplicates. C. The indistinguishable enrichment profiles of the neo-X (red) and neo-Y (blue) for the two chromatin marks are plotted with Loess smoothing curves overlaid. Correlations of the neo-X and neo-Y enrichments at individual windows can be found in Supplementary Figure 6.

**Table 2.** Differential expression between neo-X and neo-Y alleles

|  | Median log2(neo-X /neo-Y) |  | No. of genes<br>passing filter* | Significantly differentially expressed alleles** |  |  |
| --- | --- | --- | --- | --- | --- | --- |
|  | RNA-seq | DNA-seq |  | Both (% of genes) | neo-X biased (% of both) | neo-Y biased (% of both) |
| d.alb 03 (Y1) | 0.0498 | 0.0313 | 2391 | 37 (1.6%) | 28 (75.7%)† | 9 (24.3%) |
| d.alb OKNY-2 (Y2) | 0.0000 | 0.0123 | 2128 | 26 (1.2%) | 18 (69.2%) | 8 (30.8%) |
| d.alb KM55 (Y3) | 0.0580 | 0.0590 | 3181 | 168 (5.3%) | 115 (68.5%)† | 53 (31.5%) |

\* Genes with <5 reads in both alleles are removed
\*\* False discovery rate corrected $p < 0.05$ (Fisher's Exact Test)† Significantly more neo-X biased genes $p < 0.01$ (Binomial Exact Test)

In addition to gene expression differences, we compared heterochromatin and dosage compensation profiles of the sex chromosomes. We assayed two histone marks in male *D. albomicans* larvae using allele-specific ChIP-seq: H3K9me3, which is associated with silencing heterochromatin [44,47,48], and H4K16ac, which is found at the dosage-compensated, hypertranscribed X of male Drosophila [18]. As expected, the repressive histone modification H3K9me3 is enriched at the dot chromosome and the ends of chromosomes corresponding to pericentromeric regions (Figure 5B). The neo-sex chromosomes show no elevation in H3K9me3 enrichment relative to other chromosomes (Figure 5B), and the neo-Y and neo-X show nearly indistinguishable levels of the repressive H3K9me3 mark across the chromosome (Figure 5C), and enrichment levels across chromosome windows are highly correlated (Supplementary figure 6). Thus, global H3K9me3 levels reveal no evidence of neo-Y heterochromatinization. Similarly, we see no bias for the dosage compensation mark H4K16ac along the neo-X chromosome (Figure 5C, Supplementary figure 6), despite substantial enrichment on the old X Muller A (Figure 5B). This indicates that the neo-X has not yet evolved dosage compensation through H4K16ac, and is consistent with a lack of systematic up-regulation of the neo-X at the transcript level (Figure 5A). These results show that the neo-sex chromosomes have no clear signs of epigenetic differentiation, consistent with overall similar expression levels of the neo-X and neo-Y.

### Beginning neo-Y degeneration correlates with neo-Y haplotype age

Theory predicts that non-recombining neo-Y chromosomes should accumulate deleterious mutations, and the amount of degeneration should correspond to its age [29]. As mentioned above, we find that the different neo-Y haplotypes were formed at different time points, and stopped recombining between roughly 90,000 to 130,000 years ago. Despite the absence of chromosome-wide epigenetic differentiation between the neo-sex chromosomes, the number of genes with significant neo-X biased expression is more numerous than neo-Y biased genes for all three neo-Y types (Table 2, Figure 6A; note that this difference is only significant for Y3 and Y1 as they have more genes, p < 0.01, Binomial Exact Test). Most interestingly, the number of neo-X biased genes is correlated with the age of the neo-Y (Table 2, Figure 5A, Figure 6A), suggesting that older neo-Y chromosomes have more genes with reduced expression relative to the neo-X. Thus, neo-X biased gene expression may reflect the beginning stages of differentiation when degeneration of the neo-Y happens on a gene-by-gene basis.

**Figure 6.**
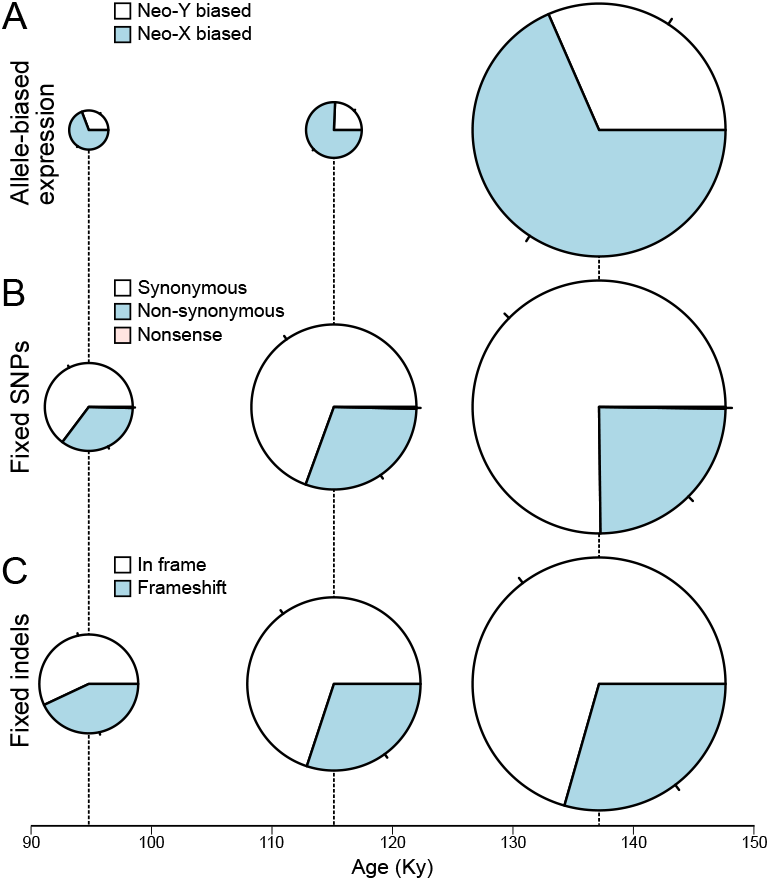
Extent of neo-Y degeneration is correlated with haplotype age. A. The proportion and number of genes with significant neo-X and neo-Y bais are plotted for each of the three haplotypes. B. Same as A, but with different classes of fixed and haplotype-specific SNPs found in CDS. C. Same as B, but with indels found in CDS. Size of pie charts are proportional to the total number of genes/mutations for each haplotype.

To determine whether older neo-Y haplotypes are associated with more deleterious DNA mutations, we focused on the region (7.6-48.6Mb) where all haplotypes differ, excluding the small haplotype found in the taiwanese line d.alb.03 (Figure 3A). We find that the number of fixed SNPs increases with the age of the haplotypes (Table 3, Figure 6B). While there are few nonsense fixations which are diagnostic of functional degeneration, the number of non-synonymous fixations correlates with age, with Y3 having the most (n = 969) and Y2 having the least (n = 478) amino-acid changes. Given few nonsense mutations, we additionally used the number of neo-Y specific indels as a proxy for functional decay of the neo-Y. Indeed, the oldest Y3 haplotype has the most fixed indels (n = 11,108), the intermediate Y1 has 5,615, and the youngest Y2 has the fewest indels (n = 3,385), but only a small fraction of indels are found within coding sequences (Table 3). Similarly, the oldest haplotype Y3 has the most number of indels that are within coding sequences (n = 163) and creates frameshift (n = 49) while the youngest haplotype Y2 has the fewest (Table 3, Figure 6C). We see the same trend with frameshift within 500bp of genes, some of which are likely to cause regulatory changes. Despite the small number of indels that are likely to be disruptive, these results demonstrate that the extent of degeneration in these young neo-Ys is correlated with their age. Curiously, haplotype-specific indels and SNPs appear to be unevenly accumulated across the chromosome (Supplementary Figure 7). Despite having fewer indels for most of the chromosome, Y1 appears to have similar numbers to that of Y3 up to ~15Mb. Additionally, Y2 has a spike of indels between 27-29 Mb which may represent a larger structural variant.

**Table 3.** Degeneration of neo-Y haplotypes

|  | No. of substitutions (7.6 - 48.6Mb) |  |  |  |  | No. of indels (7.6 - 48.6Mb) |  |  |
| --- | --- | --- | --- | --- | --- | --- | --- | --- |
|  | All | Within<br>CDS | Syn. | Non-syn. | Non-<br>sense | All | Within<br>CDS | Frame<br>shifts |
| Y1 | 13274 | 2572 | 1786 | 777 | 9 | 5615 | 163 | 49 |
| Y2 | 7480 | 1370 | 887 | 478 | 5 | 3385 | 93 | 40 |
| Y3 | 20263 | 3942 | 2694 | 969 | 9 | 11108 | 238 | 70 |

### Independent and gene-by-gene downregulation of neo-Y alleles

Interestingly, zero neo-X biased gene is shared across all three neo-Ys and no more than four are shared between any two Y-types (Figure 7A), arguing that they are independently degenerating. In addition, the lack of shared loci of neo-sex chromosome differentiation suggests that degeneration began only after the establishment of the locked neo-Y haplotypes when male recombination ceased. Furthermore, we find that across all neo-Ys, the neo-X biased genes are significantly depleted of testes-biased genes and enriched for somatic genes (Figure 7B). To ensure that the paucity of testes-biased genes is not a technical artifact of using RNA-seq from head samples, we identified neo-X-biased genes in the testes of KM55 and similarly found significant depletion. Despite independent degeneration of the neo-Y types, genes with male-specific function are thus likely to be shielded from the initial accumulation of deleterious alleles.

**Figure 7.**
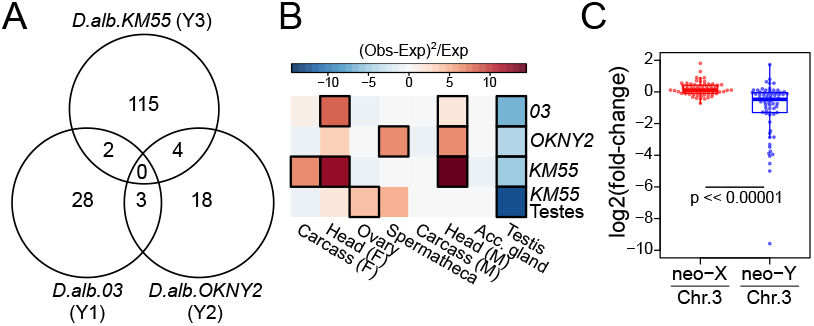
Independent gene-by-gene downregulation of neo-Y alleles. A. Venn diagram illustrates the number of neo-X biased genes that overlap between the neo-Y haplotypes. B. The heatmap depicts over- and under-representation of tissue-bias among neo-X-biased genes among the different neo-Ys. Colors represents the Chi-square statistics, modified to include enrichment (+) or depletion (-); significant cells are boxed with black borders (p < 0.05, see materials and methods). C. For genes with neo-X bias in strain *D.alb.03,* we determined their allele-specific expression in reciprocal hybrid males with Chr. 3 and either neo-X or neo-Y. The distribution (boxplot) and the gene-by-gene (points) fold-difference of the alleles are plotted in log scale. While the neo-X and Chr. 3 alleles have similar expression, the neo-Y vs. Chr. 3 fold difference in significantly lower (Wilcoxon Rank Sum Test, p << 0.00001).

To determine whether the neo-X bias is due to up-regulation of the neo-X alleles or down-regulation of the neo-Y alleles, we conducted reciprocal crosses producing F1 hybrids between *D. albomicans* and *D. nasuta;* sons sired by *D. albomicans* have neo-Y and Chr. 3 while sons sired by *D. nasuta* have neo-X and Chr. 3, thus allowing us to determine expression differences that are due to cis-regulatory divergence between the neo-X, neo-Y, and the ancestral Chr. 3 [49]. Down-regulated neo-Y alleles will have lower expression than the neo-X and, importantly, Chr. 3 alleles. For the genes with neo-X bias, the neo-X and Chr. 3 alleles have similar expression levels, but the expression of neo-Y alleles are on average half (mean = 0.48) of their Chr.3 counterparts (Figure 7C). These results reveal that neo-Y down-regulation, not neo-X up-regulation, primarily accounts for the neo-X bias, consistent with gene-by-gene degeneration of the neo-Y alleles.

## DISCUSSION

### Multi-step model for generating multiple neo-Y haplotypes with single fusion

Neo-Y chromosomes of Drosophila have served as prominent models to study the evolutionary and molecular processes resulting in the degeneration of a non-recombining chromosome [4]. Unique to *D. albomicans,* the origination of its neo-Y chromosome occurred in an ancestor without male achiasmy, and our population genetic and phylogenetic analyses shed light on the complex evolutionary history of *D. albomican’s* neo-Y chromosome. We found that the presence of male recombination created a diverse array of neo-Y haplotypes that are now geographically distributed in different parts of the species range, and propose a multi-step model to account for our results and previous models of the karyotypic evolution of *D. albomicans* (Figure 8). Laboratory crosses between *D. albomicans* and *D. nasuta* have suggested that the fusion between Chr. 3 and the X is more likely to be established first, as opposed to a Chr. 3 – Y fusion[40]. After the first Robertsonian fusion creating the neo-X [40], the neo-X is free to recombine with Chr. 3 in a meiotic trivalent both in males and females (Figure 8A). The second fusion, forming the neo-Y, occurred between the Y and a Chr. 3 that harbored a large *D. nasuta* haplotype near the centromere. This fusion is likely under positive selection to resolve mis-segregation due to the trivalent pairing during meiosis [40]. While this produced a severe bottleneck, the presence of male recombination allowed the neo-Y to regain diversity through exchange with the neo-X, which also broke down the *D. nasuta* introgression. However, once male recombination stopped in a particular population, the diversity on the neo-Y rapidly decreases, eventually leading to the fixation of one Y haplotype in the population, due to drift and possibly positive selection for male beneficial alleles (Figure 8B). Unlike on recombining chromosomes, deleterious mutations cannot be unlinked on the neo-Y and will increase in number and frequency in the population (see introduction). Because different isolated populations would likely fix different neo-Y haplotypes, overall diversity can be maintained on the neo-Y, despite high differentiation among haplotypes (Figure 2C). Our dating of neo-Y haplotypes suggests that male recombination stopped at different times for the different Y haplotypes. This raises the possibility that the mutation causing the cessation of male recombination in *D. albomicans* spread sequentially from mainland Asia through the islands.

**Figure 8.**
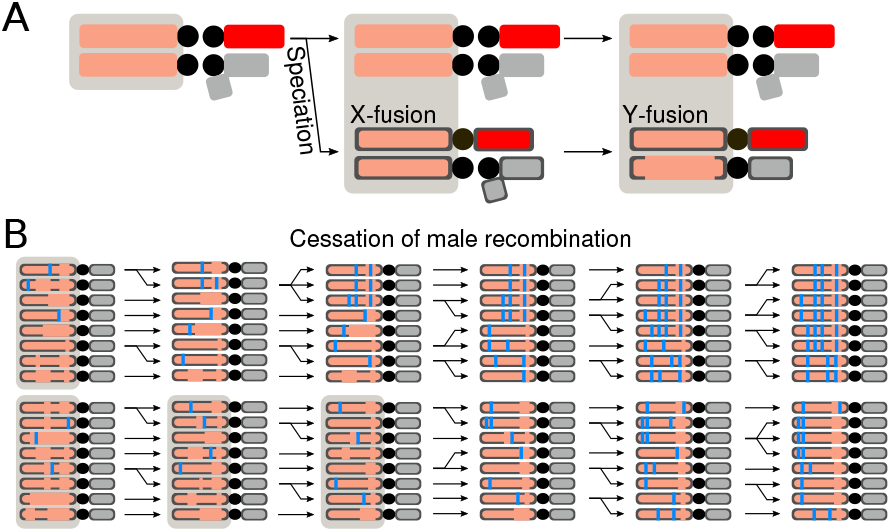
Multi-step model for neo-Y evolution. A. The sequential fusions of the chromosomes and the species split are depicted in the schematic. To differentiate the *D. nasuta* and *D. albomicans* chromosomes after the species split, the latter has black outlines. Chromosomes that can recombine with each other are encompassed by gray boxes. Following the neo-X fusion, the neo-Y emerged via fusion with a chromosome that harbors a block of Chr. 3 introgression. B. Prior to the cessation of male recombination, the diversity of neo-Y comes from primarily from recombination with the neo-X and Chr. 3. During this time, the Chr. 3 introgression on the neo-Y is broken down by recombination with the neo-X. After, the diversity is rapidly lost leading to the fixation of different haplotypes in different populations. Deleterious mutations (blue bars) begin to accumulate on the neo-Y after the cessation of recombination. The population in which male recombination stopped earlier is expected to have more degenerate neo-Ys.

All sampled neo-Ys of *D. albomicans* show the remnant of a *D. nasuta* introgression near the centromere, raising the question of why the *D. nasuta* segment was not completely lost while the neo-Y could still recombine. The introgressed segment could have been maintained randomly, or it could contain male-beneficial alleles and represent an adaptive introgression [50,51]. Another possibility is that the introgressed region resides close to the pericentromeric heterochromatin where recombination is suppressed; however, the heterochromatic mark H3K9me3 does not appear to be enriched in the segment. Preservation of the introgressed region could also be due to structural differences between the neo-sex chromosomes which prevented ancestral recombination between the neo-X and neo-Y. Indeed, in an experimental cross between *D. nasuta* and *D. albomicans,* no recombinants were recovered within this region [52]. Thus, we speculate that the neo-X harbors an inversion around this region that is absent on the neo-Y and Chr. 3, thereby preventing ancestral recombination between the neo-X and neo-Y and preserving the *D. nasuta* introgression.

### Cessation of male recombination initiates degeneration

The neo-X and neo-Y of *D. albomicans* are among the youngest sex chromosomes investigated to date. Overall, we find no global differences in levels of expression between neo-sex linked genes, and nearly indistinguishable heterochromatin and dosage compensation profiles between the neo-X and neo-Y. However, expression differences at individual genes and biased accumulation of indels suggests that the neo-Y chromosomes are showing early signs of degeneration, and the amount of differentiation is correlated with the Y ages. In particular, we identified four neo-Y haplotypes of different age that are geographically distributed. The oldest Y3 haplotype has the highest number of fixed nonsynonymous changes, indels, neo-X biased genes, indicating the most extensive degeneration; the younger Y1 and Y2 groups show less extensive degeneration. The paucity of overlapping genes that are neo-X biased between the haplotypes suggest that the degeneration began only after the haplotype became locked when males stopped recombining. We show that neo-X biased expression is primarily due to down-regulation of the neo-Y alleles, as revealed by comparison with D. nasuta’s Chr. 3 in hybrids. At this early stage of sex chromosome differentiation, down regulation of neo-Y genes is likely to be on a gene-by-gene basis, and we indeed find several genes with significantly higher levels of neo-X expression in all three Y groups (Table 2), which have also fixed indels that cause frameshifts. Unlike the typical trajectory of neo-Y in Drosophila where the male-exclusive transmission results in the immediate cessation of recombination, meiotic exchange prior to the halt of male recombination maintained the diversity of the chromosome and likely prevented the accumulation of deleterious alleles. While the neo-X and neo-Y formed shortly after the species split from *D. nasuta* [35] (Figure 4G), depending on when male recombination was abolished the age of a neo-Y haplotype will appear younger and less degenerate than expected based on when the neo-Y fusion occurred.

While the differentiation of the neo-Y is atypical compared to other Drosophila, it is similar to Y formation in organisms with male recombination. The neo-Y chromosome of the Japan Sea Stickleback (Gasterosteus nipponicus) which formed via a Y-autosome fusion shows little signs of degeneration outside of the non-recombining region [53]. Similarly, the pseudoautosomal regions on the mammalian Y (PAR-Y) chromosome retains the ability to pair and recombine with the X counterpart, despite having diverged over 180 million years ago [54–56]. Although the bulk of the chromosome is highly degenerate, recombination allows the PAR-Y to maintain genes important for development and cellular function [57]. Outside of the PAR-Y recombination with the X became suppressed due to multiple large scale inversions [58,59]. Our study of the neo-Y formation in *D. albomicans* offers a unique look at the transition of recombination suppression and provides insight on the rate of degeneration immediately after this transition.

### Geographically restricted distribution of neo-Y haplotypes that are independently degenerating

After the cessation of recombination, diversity is expected to rapidly diminish due to a combination of Hill-Robertson effect, Muller’s ratchet, and selective sweeps. Indeed, our results revealed that individuals within a geographically isolated region have nearly identical neo-Ys belonging to the same haplotypes. While more sampling of neo-Ys is required to fully capture the geographical distribution of different haplotypes, our results nonetheless paints an intriguing picture of independent fixation of different haplotypes in different isolated populations followed by independent initiation of degeneration. Given that the different haplotypes have different ages, the cessation of recombination likely spread from the oldest population from Indochina (Y3), to Taiwan (Y1), to Okinawa (Y2). The mainland Japan population appears to be a recent invasion from the Taiwanese population [34], supporting the similarity between the Y1 and Taiwanese haplotypes. Under this model, the sequential spread of the allele(s) to suppress male recombination is likely aided by positive selection, as the fixation has to take place in each population to result in the species-wide fixation as currently observed [43].

Our results also show that the neo-Y groups share very few indels and down-regulated neo-Y alleles, indicative of independent degeneration. However, the small sets of genes undergoing degeneration, remarkably, show a similar attribute – they are all depleted for genes with testes-biased expression. This is consistent with findings that male-biased genes are maintained for longer periods of time on the degenerating neo-Y-chromosome of *D. miranda,* likely due to selection to preserve those important for male-exclusive function [60]. Thus, while the independent degeneration of multiple neo-Y haplotypes may be unique given the unusual evolutionary history of *D. albomicans,* the pattern of degeneration follows a typical and generalizable trajectory.

## MATERIALS AND METHODS

### Genotyping samples from whole genome sequences

See Supplementary table 1 for all strain and sequence information. All samples were mapped with bwa (v.0.7.15) mem [61] to the reference under default settings for pair-end reads. We then processed the mapped reads according to GATK best practices which includes removing duplicates with MarkDuplicates in Picard tools (v.2.18.14) and sorting and merging with Samtools (v.1.5) [62,63]. We genotyped each chromosome arms separately using GATK’s (v3.8.0) HaplotypeCaller under default parameters. Note, because of a known but unaddressed bug in GATK where it would randomly crash, we ran HaplotypeCaller in 5 Mb windows (using the -L setting) for each chromosome to avoid restarting the entire run every time the program crashes. The genotype files (.vcf) are then merged together after all calls are complete.

Depending on the samples required for the analyses, we first filtered out the samples that are unnecessarily, followed by removal of the indel sites and sites with missing genotypes. We then filtered for a minimum coverage of 5x and genotype quality of 20. For example, for the phylogenies with only the neo-X and neo-Y, we kept only *D. albomicans* samples and the one outgroup before subsequent filtering. This is to ensure that we are maximizing the number of sites we keep for each analysis, since the probability of one sample failing to pass the filters, which will cause the site to be removed, increases with more samples. All filtering was accomplished using bcftools (v.2.26.0) [64].

### Inferring and validating the neo-Y genotypes

For each strain, we generated a vcf file containing both the female and male genotypes from which we identified sites that are homozygous in the female but heterozygous in the male. If one of the heterozygous alleles in the male is the same as the female allele, the neo-Y genotype is then the other allele. Sites that are heterozygous in the females are left as missing in the neo-Y.

To determine the efficacy of this computational pipeline, we identified HindIII cut sites on the neo-X of the reference strain by grepping for the restriction sequence AAGCTT then selecting those that are disrupted by neo-Y SNPs. Primers flanking the cut sites were designed using NCBI’s Primer-BLAST (Supplementary table 2) [65] and ordered from IDT. PCR amplicons from female and male gDNA were digested with HindIII overnight at 37 °C. Females, homozygous for the cut sites, are expected to have fewer bands than the males which are heterozygous for the cut and uncut alleles (Supplementary table 2).

For Y-linked indels, we used the same strategy, but also included male-specific homozygous indels as Y-linked. When multiple indels are within 50bp of each other, only the first one is kept to avoid mapping errors near indels. Frameshifts are categorized as indels within CDSs that are not multiples of 3.

### Population genetic estimates

Both *F_ST_* and *π* were estimated using vcftools (v.0.1.15) with the parameters --window-pi and --fst-window-size in 50kb windows [63]. *D_XY_* was estimated using the popgenwindows.py from the genomics_general package by Simon Martin (https://github.com/simonhmartin/genomics_general). Number and proportion of shared polymorphisms between the neo-X and neo-Y were calculated using a custom perl script.

### Generating strain-specific references

We isolated each sample (or neo-X/neo-Y) into its own vcf and kept only non-reference sites. Using GATK’s FastaAlternateReferenceMaker we generated a “pseudo-reference” fasta allowing for IUPAC ambiguities. The names of the chromosomes are then modified to include the strain information.

### Phylogenetic Analyses

The individual pseudo-references are divided into 200kb windows using samtools faidx and concatenated into one fasta for each window. Maximum likelihood trees were generated using RAxML (v.8.2.11) [66] with the command: raxmlHPC-PTHREADS-AVX -f a -x 1255 -p 555 -# 100 -m GTRGAMMA -s input.fa -n output.tree -o root_species.

The resulting trees were visualized using the program Densitree [67] or plotted in R with the package Phytools [68]. To evaluate tree topologies, we used the R packages ape and phytools [68,69], in which the is.monophyletic function was used to determine whether the neo-Ys are monophyletic.

### Assigning haplotypes to individual neo-Ys

For each 200kb window trees, the number of clades that contain exclusively (monophyletic) neo-Ys are first determined. Neo-Ys that are in the same monophyletic clade are then assigned the same haplotype, and different haplotypes are given a different color in the diagram. To avoid interruption of haplotypes by erroneous trees, any solo window flanked by a different topology on both side is assigned matching topology to the neighbors. For the age estimates, branch lengths are taken between nodes with phytools. When there are multiple individuals in the sister branch for the *s*-to-tip lengths, the lengths are averaged.

### Allele-specific RNA-seq and ChlP-seq

RNA were extracted from 10 male heads with Trizol, and libraries were prepared using the TruSeq Stranded Total RNA kit (Illumina). Pair-end reads are first aligned to the reference under default settings with bwa mem. Reads aligning to the neo-X which includes both neo-X and neo-Y reads are then extracted using samtools, and converted back into fastq with the bamtofastq function in Picard Tools. The fastq reads are then mapped to an index containing both the strain-specific neo-X and neo-Y sequences, and only uniquely mapping reads are kept. The read counts are then tallied using featureCounts in the Subread (v.1.6.2) package [70], with an annotation file (.gtf) where the chromosome names are modified to match the names of the strain-specific neo-X and neo-Y sequences. Genes with fewer than 5 reads in both alleles are removed from our analysis. The read counts of DNA-seq samples were processed identically. For the corrected fold difference, the fold-difference from the RNA-seq data are divided by the fold-difference from the DNA-seq data. We tested the efficacy of this approach by simulating neo-X and neo-Y reads from D.alb.KM55 at 10x and 5x coverage with the package ART [71], respectively, and were able to recapitulate the 2-fold difference at the gene level (Supplementary figure 8). Reciprocal hybrids were generated by mating 5-day old virgin D.alb.03 with male D.nas.00 or vice versa. The RNA-seq data were processed similarly with the addition of D.nas.00 specific reference.

For ChIP-seq, single male larvae were selected and processed following the low-input native chromatin immunoprecipitation protocol [72] with H3K9me3 and H4K16ac antibodies from Diagenode. The libraries were prepared using the SMARTer Universal Low Input DNA-seq Kit (Takara, formerly Rubicon). We mapped the reads similarly to the allele-specific RNA-seq, but used the genomeCoverageBed from bedtools (v.2.26.0) [73] to convert the uniquely mapped neo-X and neo-Y reads into read counts across the chromosome. We then extracted the sites that differ between the neo-X and neo-Ys. For both the input and the immunoprecipitation samples, the read counts at each site were then normalized by the median autosomal coverage. The enrichment at each window is the average across the sites.

### Tissue enrichment and depletion

Available tissue specific RNA-seq [74] were mapped to the reference genome and the read counts at genes are converted into RPKM. The tissue-specificity, tau, is then determined [75], and genes with tau > 0. 5 are deemed tissue biased and assigned to the two tissues with the highest expression. The overlap of tissue-biased and neo-X biased genes are then determined. To increase the power of this assay, instead of using the small number of neo-X-biased genes that pass the cut-off of multiple-testing (FDR) corrected p-value of 0.05, we used those that pass the nominal p-value of 0.01, increasing the number of neo-X-biased genes to 69 (alb.03), 61 (alb.OKNY), and 180 (alb.KM55). For any given Y-type-by-tissue overlap, the expected count is generated from 100,000 rounds of simulations that make two independent random draws without replacement: the number of tissue-biased genes from the set of all genes on Chr. 3 and the number of neo-X-biased genes from the set of genes passing filter (see table 2, column 4). The number of overlapping genes is determined from the two draws. The expected count is then the median of the distribution from the simulations and the P-value is determined from the percentile of the actual/observed counts.

